# CryoEM structure of the Nipah virus nucleocapsid assembly

**DOI:** 10.1101/2020.01.20.912261

**Authors:** De-Sheng Ker, Huw T. Jenkins, Sandra J. Greive, Alfred A. Antson

## Abstract

Nipah virus is a highly pathogenic zoonotic RNA virus, causing fatal encephalitis in humans. Like other negative-strand RNA viruses including Ebola and measles, its genome is wrapped by the nucleocapsid (N) protein forming a helical assembly. Here we report the CryoEM structure of the Nipah nucleocapsid protein-RNA assembly, at near atomic resolution. The N protein wraps the RNA genome with a periodicity of six nucleotides per protomer, around the outer edge of the helical assembly, in common with other paramyxoviruses. This structure uncovers details of the nucleocapsid assembly, demonstrating the role of the N-terminal arm of the N protein in the formation of the helical assembly and revealing details of the sequence-independent coordination of RNA binding in the “3-bases-in, 3-bases-out” conformation. CryoEM analysis also reveals formation of clam-shaped assemblies of the N-protein, mediated by intersubunit interactions involving several N protein loop regions.

## Introduction

Nipah virus (NiV) is an emerging zoonotic virus, causing syncytium formation, vasculitis and encephalitis, with a high mortality rate in humans^1^. It was first discovered during the outbreak in Malaysia in 1998^2^ and is one of the members of the newly delineated *Henipavirus* genus in the *Paramyxoviridae*. There is no effective vaccine against NiV and there are currently annual NiV outbreaks in Bangladesh and India for which there is no effective treatment.

As with other negative strand RNA viruses, the NiV genome is encapsidated by the nucleocapsid (N) forming a long spiral-like structure^3^. This assembly protects the viral genome from RNA degradation and the host’s intracellular defence systems. Only the encapsidated form of viral genome can serve as a template for the transcription of mRNA or nascent replication products by the viral RNA dependent RNA polymerase (RdRp)^4^. The newly synthesized viral RNA genome is encapsidated by nascently translated N protein and directed to the viral matrix protein, located in the inner side of the host cell membrane, for virus packaging via budding^5^, to complete the viral life cycle.

Several questions remain unanswered about the mechanism of viral genome encapsidation by the N protein of NiV, and more broadly, other Paramyxoviruses. It is unclear what generates specificity during recognition of the viral RNA genome by the N protein and which factors facilitate the selection of the resulting nucleocapsid, containing the viral genome, for virion packaging. Although recombinant expression of the NiV N protein in bacteria^6^, yeast^7^ and insect cells^8^ leads to the self-assembly of nucleocapsid-like structures containing cellular RNA, no structural data on the NiV N protein assembly are so far available. Here, we report CryoEM structures for several different types of assemblies formed by recombinantly produced NiV N protein, elucidating molecular interactions between N protein and RNA.

## Results

### Structure of the spiral assembly

A spiral assembly of the full length NiV nucleocapsid protein, bound to cellular RNA, was purified from recombinant expression in *E. coli*, and its structure was determined by CryoEM single-particle 3D reconstruction. 2D and 3D classification showed that the majority of the particles represent a spiral assembly comprised of 13 subunits per turn, with minor populations of particles representing a longer spiral with multiple turns, and a clam-shaped face-to-face assembly of two short spirals^9^ (Figure S1). Reconstruction of the spiral assembly (65 % of the particles) with a mask corresponding to a 13-subunit spiral turn and by imposing local symmetry, resulted in a 3.6 Å resolution CryoEM map (Figure S1). Angular distribution analysis demonstrated that while there was a preferred orientation for the particles (viewed along the central axis of the helical assembly), there was a good distribution of particles across other orientations, including side-views (Figure S1E). The final model shows that 13 nucleocapsid monomers bind to the single stranded RNA forming a left-handed spiral turn (Figure 1). Each N protein monomer is comprised of two main globular N-terminal and C-terminal Ncore (<u>N</u>ucleocapsid core) domains (Figure 1B). In addition, there are two subdomains, the N-terminal arm (NT-arm) and the C-terminal arm (CT-arm), as well as a highly disordered C-terminal tail (Ntail; <u>N</u>ucleocapsid tail)^10^ which remained unresolved in the final model.

**Figure 1:**
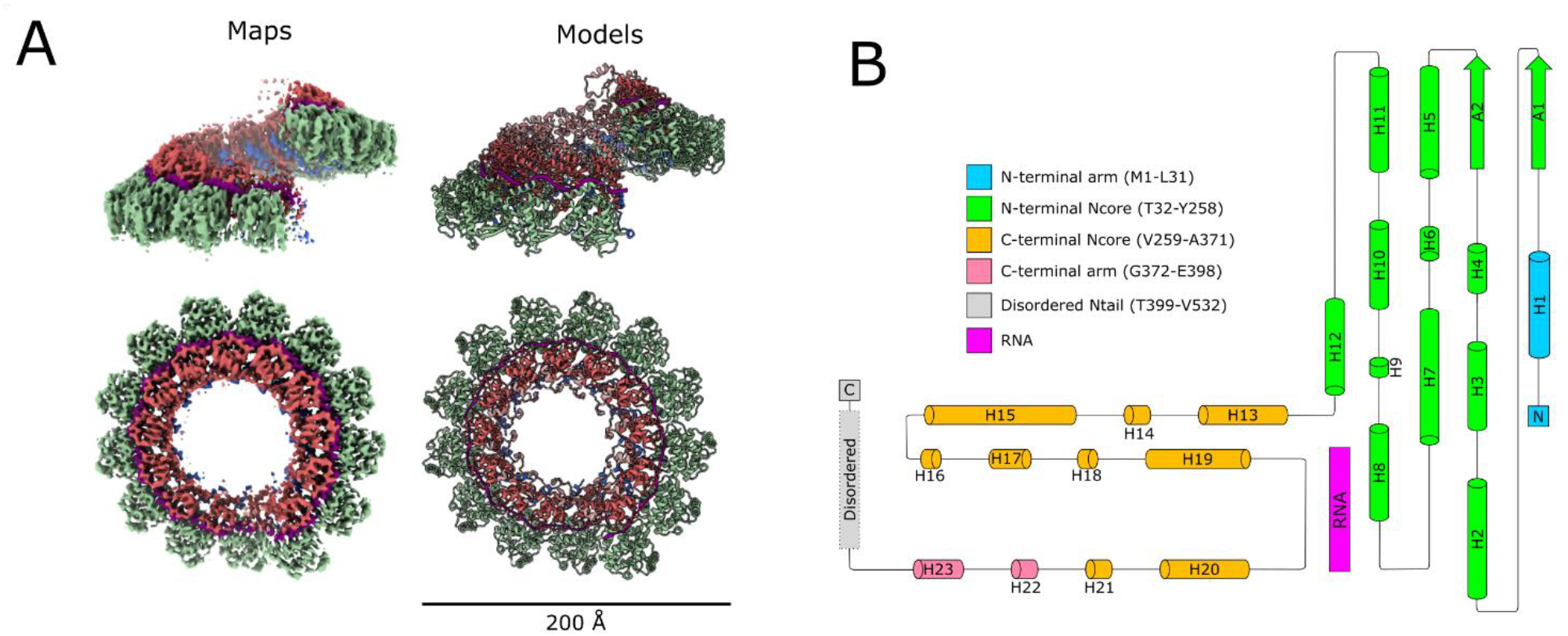
Structure of the NiV nucleocapsid protein-RNA complex. (A) CryoEM structure of the nucleocapsid complex, determined at 3.6 Å resolution. Two orthogonal views of the CryoEM map (left) are shown next to the corresponding molecular models (right). (B) A topology diagram of the NiV N protein highlighting secondary structure elements and the RNA.

### Protomer-protomer interactions within the spiral assembly

The spiral assembly of the NiV nucleocapsid is primarily formed through lateral contacts over a calculated interface area of 2998 Å^2^ between two adjacent protomers (Table S2). An aromatic residue (F11) from one protomer interacts with a triad of aromatic residues (F267, F268, Y301) within the hydrophobic groove of the adjacent protomer (Figure 2A). All of these aromatic residues are well conserved in the *Paramyxoviridae* facilitating similar protomer-protomer interactions across all family members^11^. The NT and CT arms, which have been reported to play a role in the spiral assembly of nucleocapsid^12,13^, occupy a hydrophobic groove in the C-terminal Ncore domain of the adjacent protomer. In the crystal structure of the NiV N protein monomer (pdb:4co6), a similar hydrophobic groove is occupied by a 50 amino acid segment of the NiV phosphoprotein (P) which is essential in maintaining the N protein in its RNA-free, monomeric state (Figure 2B)^12^.

**Figure 2:**
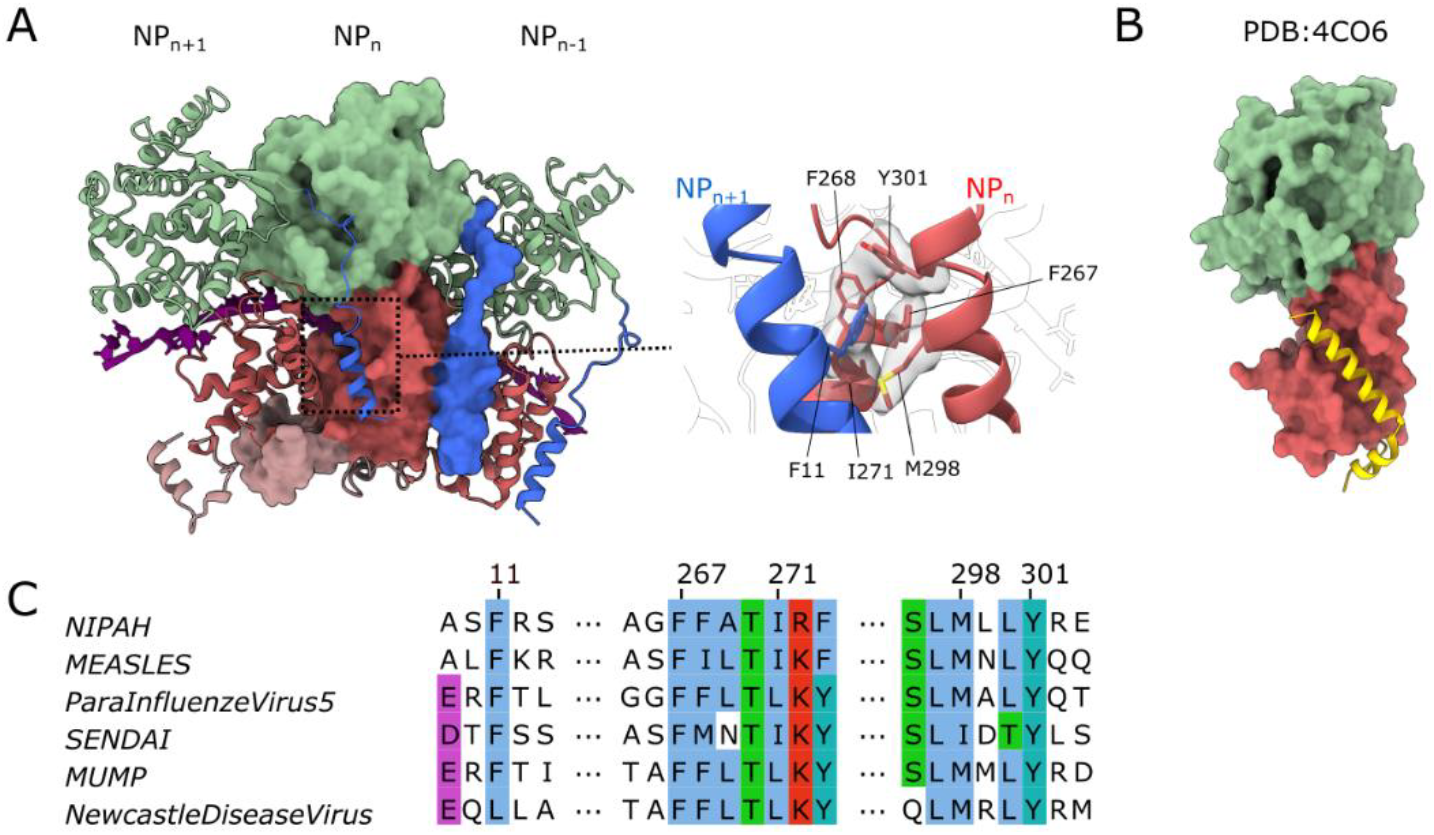
Protomer-protomer interactions within the NiV nucleocapsid assembly. (A) Three adjacent protomers, where the two outside protomers are presented as ribbons and the central protomer is shown in surface representation. A magnified view of the molecular interaction of the N-terminal arm and the C-terminal Ncore domain is shown (right) with interacting residues (sticks) displayed within CryoEM map. (B) Structure of the monomeric NiV N protein bound to the P protein (pdb:4co6)^12^ shown in the same view as the central subunit in (A). (C) Alignment of interacting residue segments, shown in the magnified view in (A), for N proteins from several Paramyxoviruses, with conserved residues highlighted using the ClustalX colour scheme.

### Protein-RNA interactions

The CryoEM map shows clear density for the single stranded RNA, modelled as a poly-uracil chain, wrapped around the nucleocapsid. The RNA molecule is bound to the protein in the classical “3-base-in, 3-base-out” conformation^11^, where the RNA chain twists 180° every 3 nucleotides, where three consecutive nucleotides with bases facing the protein are followed by 3 nucleotides with exposed bases. The structure shows that the nucleic acid lies within the charged cleft of the N protein, at the interface between the N-terminal Ncore and the C-terminal Ncore domains, in a grove lined by residue segments K178-Q200 and S344-Y354, at the outer edge of the spiral assembly (Figure 3). Within the RNA binding cleft, a series of basic (K178, R192, R193, R352) and polar (T181, Q319, S344) residues, with well-defined density, are within hydrogen-bonding distance from the sugar-phosphate backbone of RNA. Residues Q199 and Q200 from helix H8 are also have well-defined density and are oriented toward the RNA bases. These two amino acids are conserved in other members of the *Paramyxoviridae*, and as seen from structural data for the measles virus (MeV) and parainfluenza virus 5 (PIV5), make similar interactions with RNA bases^14,15^. At the interface between the two protomers, aromatic residues Y258 and Y354, one each from adjacent protomer, are positioned in close proximity to the RNA chain (Figure 3C), facilitating the twist in its conformation. This twist in the sugar-phosphate backbone of the RNA strand is assisted by a series of additional protein-RNA interactions contributed by other polar residues exposed within the RNA binding cleft. The second twist in the RNA conformation, spaced by three nucleotides from the first one, occurs in a cleft within a single protomer and is facilitated by steric hindrance from the side chain of L348 (Figure 3D). The majority of residues interacting with the RNA within the RNA-binding cleft are highly conserved among *Paramyxovirus* N proteins (Figure 3E) indicating a similar mechanism for RNA coordination.

**Figure 3.**
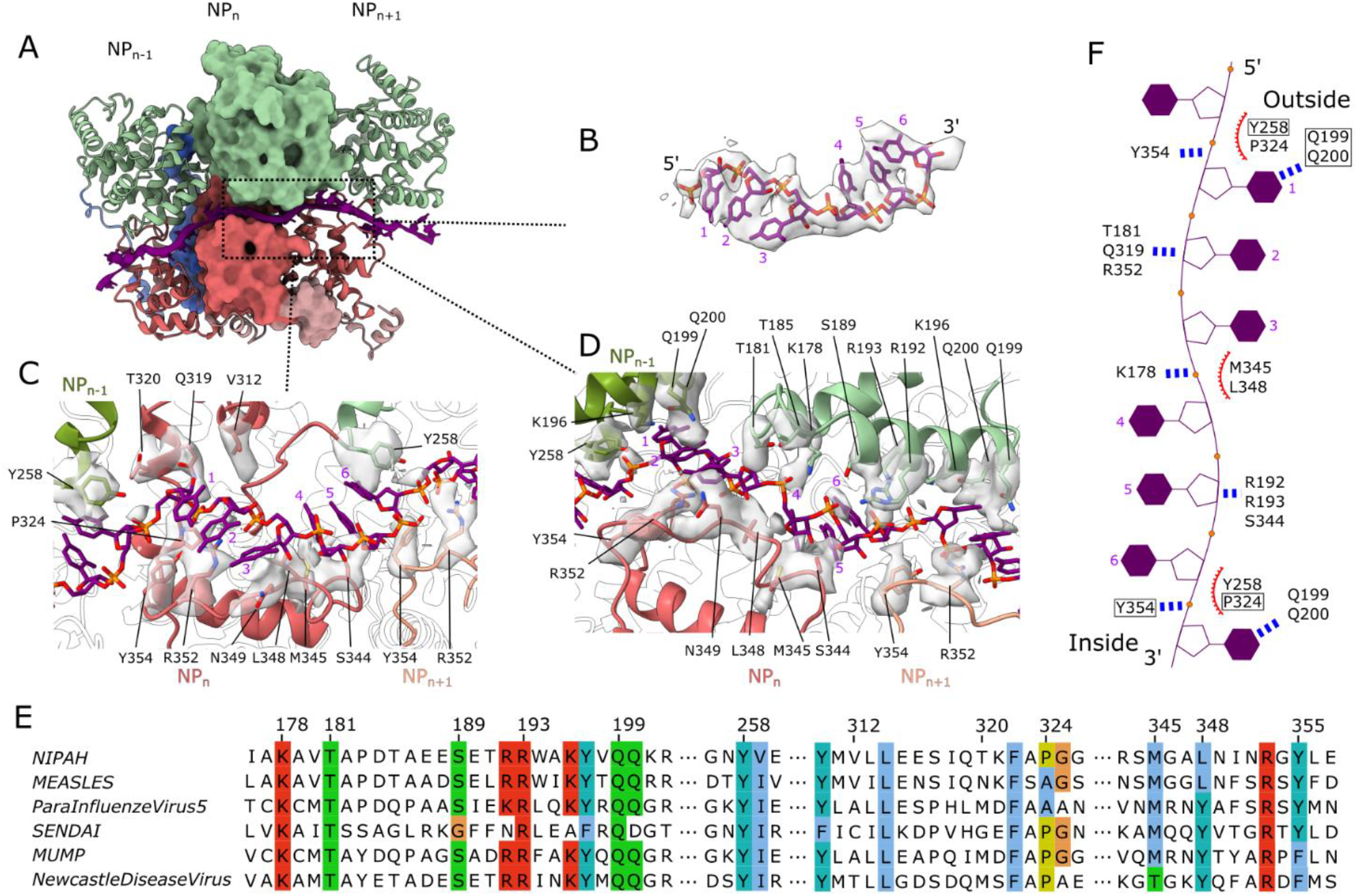
Protein-RNA interactions. (A) Three adjacent protomers shown as in Figure 2A, with RNA (purple) shown in ribbon and sticks. (B) CryoEM map corresponding to the RNA, with a fitted poly-U RNA model (sticks). (C-D) Two different views at the protein-RNA interface, detailing protein-RNA interactions. CryoEM density corresponding to the side chain atoms of interacting residues, is shown with a 3 Å distance cut-off. (E) Alignment of RNA-binding residue segments from several Paramyxoviral N proteins with conserved residues highlighted using the ClustalX colour scheme. (F) Schematic of the ssRNA conformation in complex with the N protein. The residues in close proximity to RNA are labelled. Boxed residues indicate those from the neighbouring protomer; the blue dotted lines indicate putative hydrogen bonding interactions; red curves indicate putative hydrophobic interactions.

Several residues connecting the two RNA binding segments (K178-Q200 and S344-Y354) with the rest of the N-protein are poorly defined. It is likely that the flexible nature of these regions serves to provide plasticity to accommodate and interact, in a non-sequence specific manner, with the varying sequence along the entire length of the RNA strand. The flexibility may also allow the RdRp to access the ssRNA bound to the nucleocapsid, for RNA synthesis.

In the crystal structure of the RNA-free monomeric NiV N protein, the flexible loop, residues A180-R192, was mostly disordered and positioned such that it would block access to the RNA binding cleft, suggesting that this loop needs to move out of the cleft to permit RNA binding. As seen from structure comparison, RNA binding is also accompanied by an approximately 28° rotation of the N-terminal and C-terminal Ncore domains towards each other, around a hinge region formed by the H12-H13 loop^12^, H15-H16 loop and helix H17 (Figure S2, Table S3). Similar conformational changes have also been observed for the nucleocapsid of MeV^11^.

### Clam-shaped assemblies

Aside from the common spiral assembly, about 35 % of N protein particles were found as clam-shaped assemblies which can be further classified into two primary distinct conformations, a spiral clam-shaped assembly and a semi-spiral clam-shaped assembly. The spiral clam-shaped assembly is composed of two N protein spirals stacked face to face, as seen for the Newcastle Diseases virus (NDV) N protein assembly^9^. In contrast, the semi-spiral clam-shaped assembly features one 14 protomer N protein ring and one N protein spiral stacked together (Figure S3). Asymmetric reconstruction of both assemblies leads to 4.3 Å and 4.9 Å resolution maps, respectively, with the seam region, the area between the turns of the adjacent helical assemblies, remaining poorly resolved. Models for both types of the spiral assembly were built by rigid-body fitting and real space refinement of the N protein protomer taken from the protein-RNA complex described above. The structure of the N protein monomer within these clam-shaped assemblies remains largely the same as in the spiral assembly, with the overall RMSD of 0.9 Å calculated over Cα atoms.

For both assemblies, there is a significant surface area buried at the interface between the two halves of the clam shell, with 666 Å^2^ of buried area per monomer. Interactions across this interface are mediated by hydrogen bonding and polar interactions made by loop segments A1-H2, A2-H5 and H6-H7 (Figure 4B) from each opposing protomer. The surface area, buried at the clam-shell interface of each monomer, is about five times larger than seen in the clam-shaped assembly of NDV, where only one protein loop (residues 104-124) is involved in the interaction^9^. This is due to a significantly tighter interaction between the halves of the NiV clam shell as compared to the NDV assembly. Interestingly, although sequences of these clam-shell interface loops are not conserved between NiV and NDV, or other members of the *Paramyxoviridae*, these loops are rich in glycine and proline residues (Figure 4C).

**Figure 4.**
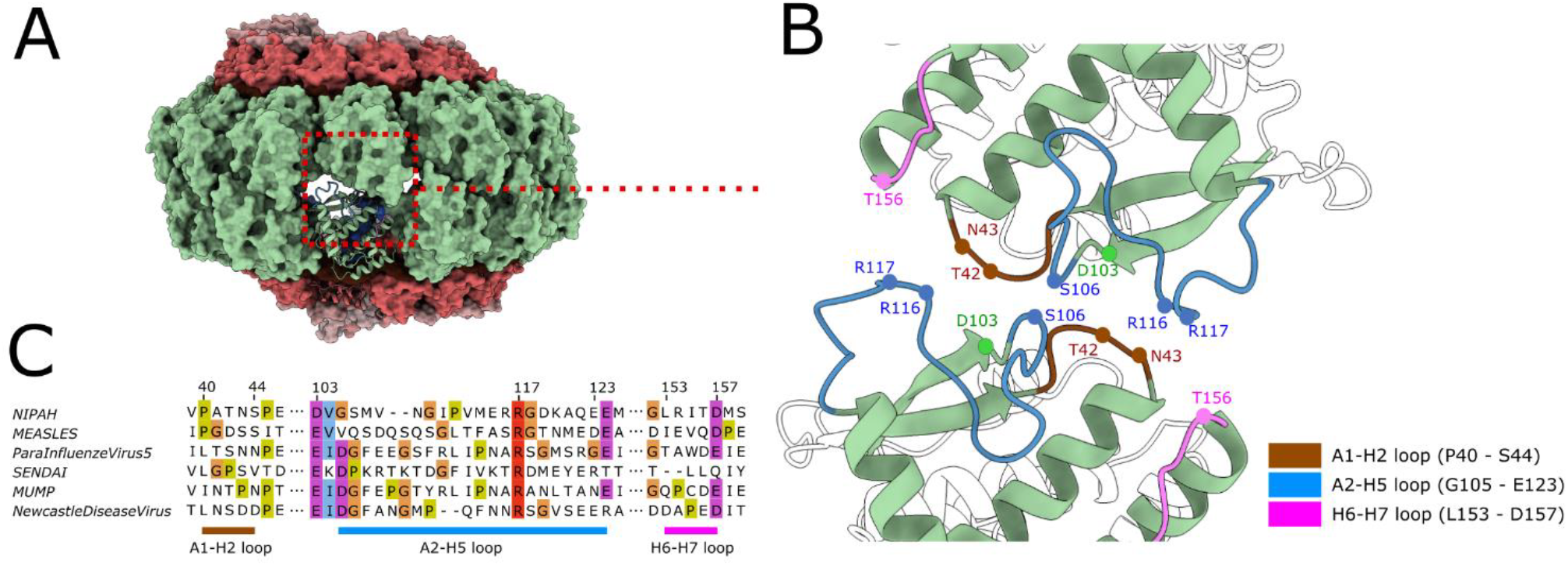
Clam-shaped nucleocapsid assembly. (A) Model of the clam-shaped nucleocapsid assembly, presented as a molecular surface, showing the interaction between two opposing, top and bottom N protein spirals, with one of the protomers shown as ribbon. (B) The clam-shaped interaction is primarily mediated by protein loops from the N-terminal Ncore domain. Putative residues involved in the interaction between the two halves of the shell are indicated. (C) Alignment of the three interacting loop regions highlighted in (B) for N proteins from several Paramyxoviruses, with glycine, proline and surrounding conserved residues highlighted using the ClustalX colour scheme.

## Discussion

RNA viral genome encapsidation by the N protein is an essential step in the life cycle of all negative-strand RNA viruses. The encapsidated genome serves as a template for both mRNA transcription and viral genome replication. Structural analysis of a spiral assembly, formed by a full-length recombinant NiV N protein, reported here (Figure 1), revealed an architecture common to other RNA-bound *Paramyxoviridae* family N proteins^9,11,14^, indicating that the overall mechanism for encapsidation of the viral genome is universal among members of this family. Upon binding to the RNA, the monomeric NiV N protein undergoes a conformational change involving a 28° rotation via the hinge motion between the two Ncore domains, effectively closing the two lobes around the RNA resulting in tight binding. The formation of the spiral assembly of the N protein is stabilised by the projection of the NT- and CT-arms onto the adjacent protomers (Figure 2).

In the protein-RNA complex, there are six single stranded RNA nucleotides bound per protein protomer, in a classical “3-base-in, 3-base-out” conformation, involving a 180° rotation, or ‘switch’, in the RNA backbone, every three bases. Such changes in the RNA conformation are facilitated by a combination of hydrogen bonding interactions and a steric hindrance with residues in the RNA binding cleft (Figure 3). In particular, the highly conserved bulky aromatic residues Y258 and Y354 found at the interface of the protomers, appear to be particularly important for RNA binding and facilitating the twist in its conformation. Point mutation of the corresponding Y258 residue in Sendai virus (SeV) prevented formation of the classical nucleocapsid protein assembly, highlighting the importance of this residue for RNA binding^16^.

While the majority of the NiV N protein particles form the typical spiral assembly, a significant proportion, 35 %, form a clam-shaped assembly, where the 5ʹ end of one encapsidated RNA faces the 5 ʹ end of another encapsidated RNA. A very similar type of assembly has been observed for the N protein from the NDV virus^9^, a distant homologue of NiV with only 30 % sequence identity, pointing at a potential role in the paramyxoviral life cycle for this type of assembly. Interestingly, two distinct types of clam-shaped assemblies are observed for the NiV N protein: (i) a spiral clam-shaped conformation, which is formed by two spirals stacking against each other; and (ii) a semi-spiral clam-shaped assembly, formed by the stacking of ring-like and spiral oligomers. Previous mutational analysis, in NDV and SeV, of residues corresponding to the A2-H5 loop (Figure 4B), which facilitates formation of the clam-shaped assembly, has resulted in up to 100% loss of *in vitro* replication while retaining the helical assembly capability of the N protein^9,17^. However, the molecular mechanism underlying this loss of activity is unknown, since the RdRp synthesizes the new RNA strand by initiating at the promoter encoded in the 3ʹ end of RNA template^18^, and hence is not expected to interact with the clam-shaped assembly that accommodates the 5ʹ end of the RNA template. No large conformational changes are observed between the monomers of the spiral and clam-shaped assemblies, suggesting that the equilibrium of the spiral and clam assemblies might be influenced by buffer conditions. Indeed, salt concentration and pH have been shown to alter the conformation and oligomeric state of the protein-RNA assemblies in Paramyxovirus^19^ and Tobacco Mosaic Virus (TMV)^20^.

The encapsidated viral RNA genome, despite being protected from degradation by nucleases, is still accessible to the viral RdRp, suggesting a yet to be understood mechanism that regulates encapsidation and transcription. The clam-shaped assembly observed here might be one of the factors that modulates the switch between mRNA transcription and viral genome replication. Another regulatory role may be contributed by the flexible Ntail region located at the C-terminus of the N protein. The N protein Ntail region has been shown both to be the site for P protein interaction^21,22^ and to play a role in regulating transcriptional activity^23,24^. It is likely that this region is orientated toward the outside of the spiral^25^, allowing contact with the P protein. Deciphering this molecular interaction in the future will provide new insights about the regulation of transcription and replication by members of the *Paramyxoviridae* family.

## Material and Methods

### Expression and purification of the NiV nucleocapsid spiral assembly

The NiV nucleocapsid gene, a kind gift from Wen Siang Tan at the Universiti Putra Malaysia, was cloned into the pET28a-Lic+ expression vector. N protein was expressed (at 16 °C) in *E. coli* BL21 Gold (DE3) Rosetta pLysS grown in 2 L of LB media. Cell pellets were resuspended in 50 mL of lysis buffer containing 20 mM Tris-HCl, pH 8.0, 1 M NaCl, 1 M Urea, 50 mM Imidazole, and 10 % (v/v) glycerol. Cells were lysed by sonication and the lysate clarified by centrifugation at 25,000 *g* for 30 min. The supernatant was then applied to a 5mL HisTrap FF column (GE Healthcare) which had been equilibrated with 5 column volumes (CV) of binding buffer containing 20 mM Tris-HCl, pH 8.0, 1 M NaCl, 50 mM Imidazole, and 10 % (v/v) glycerol. The column was washed with 10 CV of binding buffer containing 50 mM Imidazole, followed by 6 CV of binding buffer containing 100 mM Imidazole. The protein was eluted using a linear gradient from 100 mM to 500 mM imidazole over 20 CV. Eluted protein was concentrated and further purified by size exclusion chromatography (Superose 6, GE Healthcare) in 20 mM Tris pH 8.0, 500 mM NaCl. The protein was concentrated to 1 mg/mL, flash frozen in liquid nitrogen, and stored at −80 °C. The concentration of the N protein was determined using the Bradford Assay (Thermo Fisher Scientific).

### CryoEM sample preparation and data acquisition

The purified N protein was prepared on UltraAuFoil R1.2/1.3 gold support grids (Quantifoil). 3 µl of sample was applied to glow-discharged grids, blotted for 2 s with −10 force, and vitrified by plunging into liquid ethane using the FEI Vitrobot Mark IV at 4°C and 100 % relative humidity. Micrographs were collected at the Diamond eBIC facility on a Titan Krios microscope (FEI) operating at 300 keV and equipped with K2 camera (Gatan). Automated data collection was performed using FEI EPU software. 1879 movies with a total electron dose of ~41 e-/Å^2^ were recorded in counting mode over 11 s (40 frames) with a pixel size of 1.048 Å. The defocus range chosen for automatic collection was 0.5 to 2.1 µm.

### Image Processing

All datasets were processed in RELION 3.0^26^ unless stated otherwise. Micrographs were first motion-corrected using MotionCor2^27^. CTF parameters were estimated using CTFFIND4^28^. Autopicking was performed in RELION using references generated from manually picked particles. All the micrographs were manually inspected to ensure picking of rare views. A total of 217,522 particles were extracted and subjected to reference-free 2D classification to remove particles associated with noisy or contaminated classes. The resulting 189,662 particles were subjected to 3D classification using a map generated from the Measles N protein (EMDB:0141), low-pass filtered to 60 Å, as a reference model. The best 3D class was low-pass filtered to 30 Å and used as a reference model for a new round of 3D classification against the same initial set of particles. Reconstruction of the final spiral map was achieved by using all particles from the spiral classes (124,891) with a 13-protomer spiral turn solvent mask and imposing local symmetry. This improved the map quality enabling model building. For the local symmetry, masks around all 13 protomers were created and low-pass filtered to 15 Å. The local symmetry operators were generated from the search feature of the *relion_local_symmetry* and were applied during the 3D refinement in RELION. Subsequent per-particle CTF refinement and Bayesian polishing in RELION 3.1beta^29^ led to a final map of 3.6 Å resolution, estimated by the 0.143 FSC criterion (Figure S1D). The maps were postprocessed in RELION 3.1beta^29^ and are shown after B-factor sharpening.

The remaining 64,771 non-spiral particles were further subjected to 3D classification using the “clam-shaped” 3D class, obtained from previous 3D classification, as a reference model. Two major 3D classes, a spiral clam-shaped assembly (23,029 particles) and a semi-spiral clam assembly (18,979 particles), were selected. Subsequent per-particle CTF refinement in RELION 3.1beta, 3D refinement of these two 3D classes resulted in final maps of 4.3 Å (spiral clam-shaped assembly) and 4.9 Å (semi-spiral clam-shaped assembly) resolution, respectively.

### Model building, refinement and analysis

Atomic model building of the NiV nucleocapsid spiral assembly was performed using the previously reported crystal structure of the RNA free NiV nucleocapsid (pdb:4co6) as an initial model, which was docked as a rigid body into the 3.6 Å resolution CryoEM maps using UCSF Chimera’s “Fit in map” function^30^. The RNA nucleotides were added, and the model was adjusted manually using Coot^31^. Model refinement was performed using REFMAC5^32^, phenix.real_space_refine^33^, ISODLE^34^ and ERRASER2. Terminal residue segments 1-4 and 399 – 532 were not modelled owing to a lack of interpretable map features.

For the “semi-spiral clam”, and “spiral clam” CryoEM maps, the monomeric model of the RNA-bound NiV N protein (taken from this study) was fitted as a rigid body into the maps^30^. No protein models were fitted into the seam regions, due to the lack of interpretable map features in the CryoEM maps. Both models were refined using Refmac5^32^ and phenix.real_space_refine^33^.

Protein interfaces were analysed using the COCOMaps server^35^. Protein domain motion was analysed using the DynDom server^36^. Multiple sequence alignments were performed using Clustal Omega^37^ and visualised in JalView^38^. Figures showing protein/RNA structure were created using UCSF ChimeraX^39^.

## Acknowledgements

This project was supported by a Platinum Scholarship from the Tony Wild Fund to DSK and by a Wellcome Trust Award 206377 to AAA. We thank Wen Siang Tan (Universiti Putra Malaysia) for kindly providing the NiV Nucleocapsid gene. We thank Maria Chechik, Sam Hart (University of York), and Svetomir Tzokov (University of Sheffield) for help in CryoEM grid screening. We acknowledge Diamond Light Source for access to- and support from-the Cryo-EM facilities at the UK national electron bio-imaging centre (eBIC), proposal EM19832-12, funded by the Wellcome Trust, MRC and BBSRC. We are also grateful for computational support received from the University of York High Performance Computing service (Viking) and the Research Computing team.

## References

1. Torres-Velez, F. J. et al. Histopathologic and Immunohistochemical Characterization of Nipah Virus Infection in the Guinea Pig. Vet. Pathol. 45, 576–585 (2008).

2. Eaton, B. T., Broder, C. C., Middleton, D. & Wang, L.-F. Hendra and Nipah viruses: different and dangerous. Nat. Rev. Microbiol. 4, 23–35 (2006).

3. Chua, K. B. Nipah Virus: A Recently Emergent Deadly Paramyxovirus. Science. 288, 1432–1435 (2000).

4. Ogino, T. & Green, T. J. RNA synthesis and capping by nonsegmented negative strand RNA viral polymerases: Lessons from a prototypic virus. Front. Microbiol. 10, 1–28 (2019).

5. Ray, G., Schmitt, P. T. & Schmitt, A. P. C-Terminal DxD-Containing Sequences within Paramyxovirus Nucleocapsid Proteins Determine Matrix Protein Compatibility and Can Direct Foreign Proteins into Budding Particles. J. Virol. 90, 3650–3660 (2016).

6. Tan, W. S., Ong, S. T., Eshaghi, M., Foo, S. S. & Yusoff, K. Solubility, Immunogenicity and Physical Properties of the Nucleocapsid Protein of Nipah Virus Produced in Escherichia coli. J. Med. Virol. 73, 105–112 (2004).

7. Joseph, N. M. S., Tey, B. T., Tan, C. S., Shafee, N. & Tan, W. S. Production of long helical capsid of Nipah virus by Pichia pastoris. Process Biochem. 46, 1871–1874 (2011).

8. Eshaghi, M., Tan, W. S., Ong, S. T. & Yusoff, K. Purification and Characterization of Nipah Virus Nucleocapsid Protein Produced in Insect Cells. J. Clin. Microbiol. 43, 3172–3177 (2005).

9. Song, X. et al. Self-capping of nucleoprotein filaments protects the Newcastle disease virus genome. Elife 8, 1–19 (2019).

10. Habchi, J., Mamelli, L., Darbon, H. & Longhi, S. Structural disorder within Henipavirus nucleoprotein and phosphoprotein: From predictions to experimental assessment. PLoS One 5, (2010).

11. Gutsche, I. et al. Near-atomic cryo-EM structure of the helical measles virus nucleocapsid. Science 348, 704–707 (2015).

12. Yabukarski, F. et al. Structure of Nipah virus unassembled nucleoprotein in complex with its viral chaperone. Nat. Struct. Mol. Biol. 21, 754–759 (2014).

13. Milles, S. et al. Self-Assembly of Measles Virus Nucleocapsid-like Particles: Kinetics and RNA Sequence Dependence. Angew. Chemie - Int. Ed. 55, 9356–9360 (2016).

14. Alayyoubi, M., Leser, G. P., Kors, C. A. & Lamb, R. A. Structure of the paramyxovirus parainfluenza virus 5 nucleoprotein– RNA complex. Proc. Natl. Acad. Sci. 112, E1792–E1799 (2015).

15. Desfosses, A. et al. Assembly and cryo-EM structures of RNA-specific measles virus nucleocapsids provide mechanistic insight into paramyxoviral replication. Proc. Natl. Acad. Sci. 201816417 (2019). doi:10.1073/pnas.1816417116

16. Myers, T. M., Pieters, A. & Moyer, S. A. A highly conserved region of the Sendai virus nucleocapsid protein contributes to the NP-NP binding domain. Virology 229, 322–335 (1997).

17. Myers, T. M. & Moyer, S. A. An amino-terminal domain of the Sendai virus nucleocapsid protein is required for template function in viral RNA synthesis. J. Virol. 71, 918–924 (1997).

18. Jordan, P. C. et al. Initiation, extension, and termination of RNA synthesis by a paramyxovirus polymerase. PLOS Pathog. 14, e1006889 (2018).

19. Heggeness, M. H., Scheid, A. & Choppin, P. W. Conformation of the helical nucleocapsids of paramyxoviruses and vesicular stomatitis virus: reversible coiling and uncoiling induced by changes in salt concentration. Proc. Natl. Acad. Sci. 77, 2631–2635 (1980).

20. Kegel, W. K. & van der Schoot, P. Physical Regulation of the Self-Assembly of Tobacco Mosaic Virus Coat Protein. Biophys. J. 91, 1501–1512 (2006).

21. Habchi, J. et al. Characterization of the Interactions between the Nucleoprotein and the Phosphoprotein of Henipavirus. J. Biol. Chem. 286, 13583–13602 (2011).

22. Ranadheera, C. et al. The interaction between the Nipah virus nucleocapsid protein and phosphoprotein regulates virus replication. Sci. Rep. 8, 15994 (2018).

23. Cox, R. M., Krumm, S. A., Thakkar, V. D., Sohn, M. & Plemper, R. K. The structurally disordered paramyxovirus nucleocapsid protein tail domain is a regulator of the mRNA transcription gradient. Sci. Adv. 3, (2017).

24. Thakkar, V. D. et al. The Unstructured Paramyxovirus Nucleocapsid Protein Tail Domain Modulates Viral Pathogenesis through Regulation of Transcriptase Activity. J. Virol. 92, JVI.02064–17 (2018).

25. Kingston, R. L., Baase, W. A. & Gay, L. S. Characterization of Nucleocapsid Binding by the Measles Virus and Mumps Virus Phosphoproteins. J. Virol. 79, 10097–10097 (2005).

26. Zivanov, J. et al. New tools for automated high-resolution cryo-EM structure determination in RELION-3. Elife 7, (2018).

27. Zheng, S. Q. et al. MotionCor2: anisotropic correction of beam-induced motion for improved cryo-electron microscopy. Nat. Methods 14, 331–332 (2017).

28. Rohou, A. & Grigorieff, N. CTFFIND4: Fast and accurate defocus estimation from electron micrographs. J. Struct. Biol. 192, 216–221 (2015).

29. Zivanov, J., Nakane, T. & Scheres, S. H. W. Estimation of High-Order Aberrations and Anisotropic Magnification from Cryo-EM Datasets in RELION-3.1. bioRxiv (2019). doi:10.1101/798066

30. Pettersen, E. F. et al. UCSF Chimera-A visualization system for exploratory research and analysis. J. Comput. Chem. 25, 1605–1612 (2004).

31. Emsley, P., Lohkamp, B., Scott, W. G. & Cowtan, K. Features and development of Coot. Acta Crystallogr. Sect. D Biol. Crystallogr. 66, 486–501 (2010).

32. Murshudov, G. N. et al. REFMAC 5 for the refinement of macromolecular crystal structures. Acta Crystallogr. Sect. D Biol. Crystallogr. 67, 355–367 (2011).

33. Afonine, P. V. et al. Real-space refinement in PHENIX for cryo-EM and crystallography. Acta Crystallogr. Sect. D Struct. Biol. 74, 531–544 (2018).

34. Croll, T. I. ISOLDE : a physically realistic environment for model building into low-resolution electron-density maps. Acta Crystallogr. Sect. D Struct. Biol. 74, 519–530 (2018).

35. Vangone, A., Spinelli, R., Scarano, V., Cavallo, L. & Oliva, R. COCOMAPS: a web application to analyze and visualize contacts at the interface of biomolecular complexes. Bioinformatics 27, 2915–2916 (2011).

36. Lee, R. A., Razaz, M. & Hayward, S. The DynDom database of protein domain motions. Bioinformatics 19, 1290–1291 (2003).

37. Sievers, F. et al. Fast, scalable generation of high‐quality protein multiple sequence alignments using Clustal Omega. Mol. Syst. Biol. 7, 539 (2011).

38. Waterhouse, A. M., Procter, J. B., Martin, D. M. A., Clamp, M. & Barton, G. J. Jalview Version 2--a multiple sequence alignment editor and analysis workbench. Bioinformatics 25, 1189–1191 (2009).

39. Goddard, T. D. et al. UCSF ChimeraX: Meeting modern challenges in visualization and analysis. Protein Sci. 27, 14–25 (2018).

