## Supplementary document for "CryoEM structure of the Nipah virus nucleocapsid assembly"

### Supplementary materials

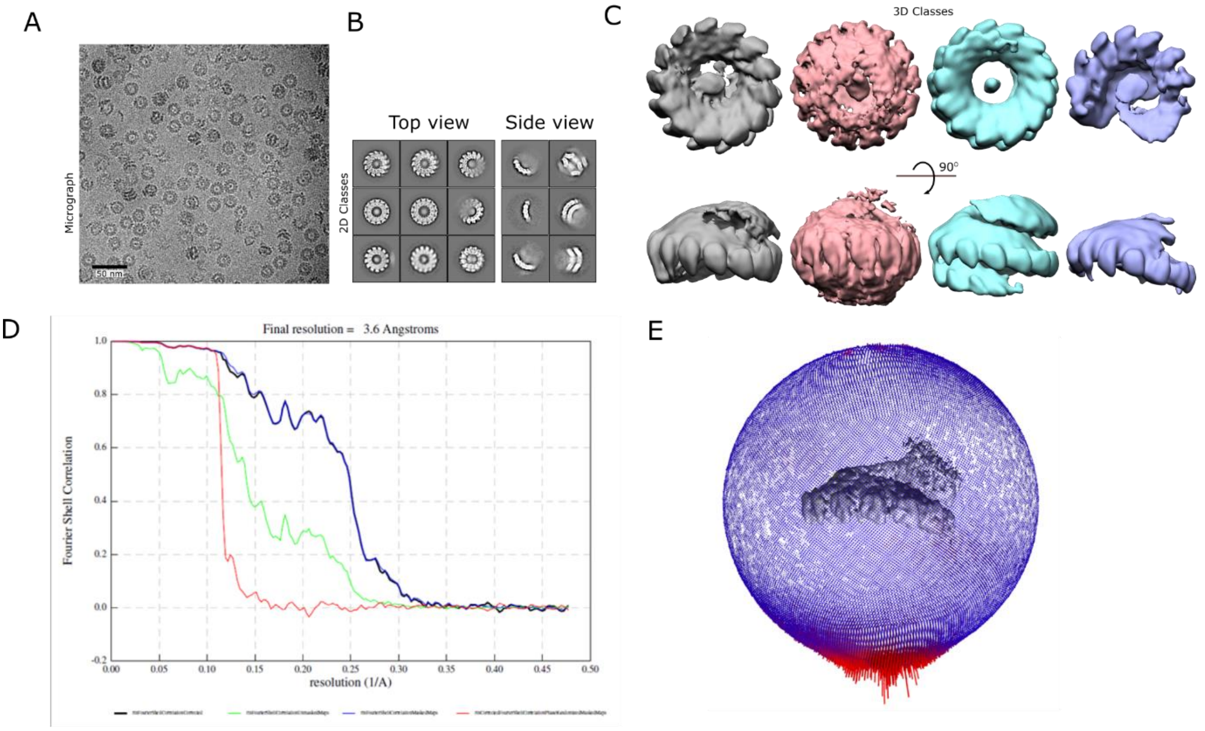


*Figure S1* ***CryoEM data processing.*** *(A) Representative micrograph of the Nipah virus (NiV) N protein spiral assembly. (B) 2D class averages of the top and side views of the N protein spiral assembly. (C) 3D class averages of the N protein spiral assembly. (D) Gold-standard FSC plot generated using RELION3.0. (E) Angular distribution plot generated by RELION3.0.*

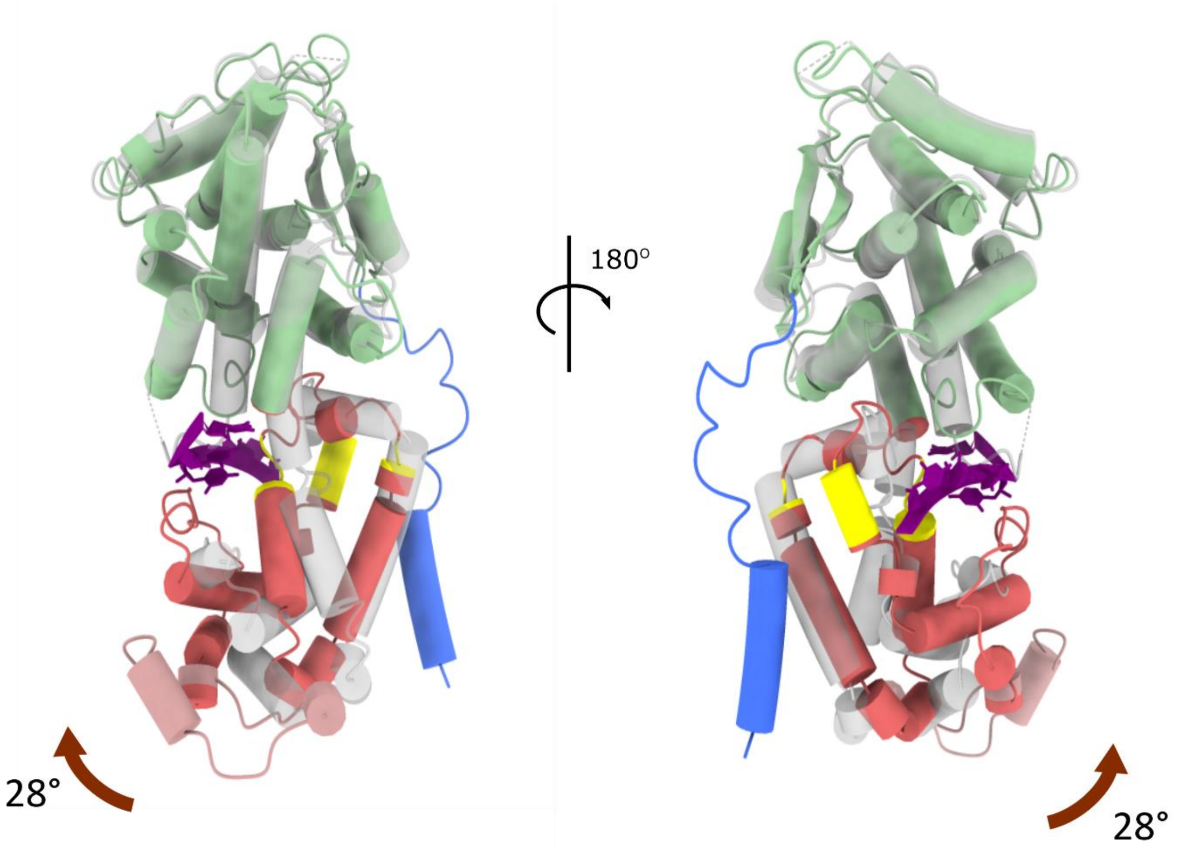


*Figure S2* ***Comparison of the Nipah N protein structure in the RNA-free and RNA-bound states.*** *Superimposed models are presented as cartoons. The RNA-free N protein (pdb:4co6)*[12] *is in semi-transparent grey; while the RNA-bound protein is in same colours as in Fig. 2A. The hinge regions are highlighted in yellow.*

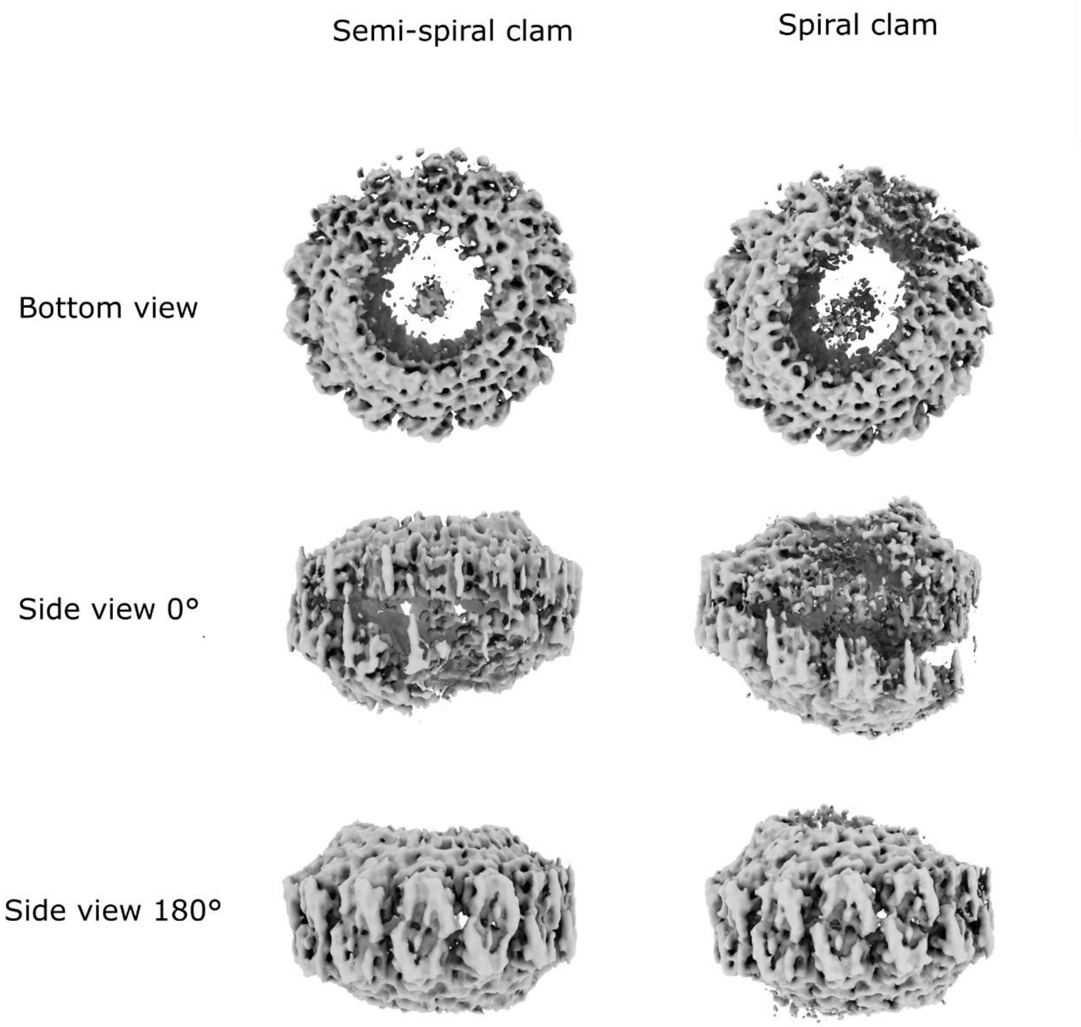


*Figure S3* ***CryoEM maps for the two major types of clam-shaped assemblies.*** *Unsharpened CryoEM maps for each of the two assemblies are shown in three different views.*

*Table S1* ***Statistics of CryoEM data collection and processing***

| | Spiral | Spiral Clam | Semi-spiral Clam |
|---|---|---|---|
| **Data collection** | | | |
| Voltage (kv) | 300 | | |
| Detector | Gatan K2 Summit | | |
| Electron exposure (e-/Å$^2$) | 41.2 | | |
| Defocus range (μm) | 0.5 to 2.1 | | |
| Pixel size (Å) | 1.048 | | |
| **Data processing** | | | |
| Symmetry imposed | C1 | C1 | C1 |
| Final particle images (no.) | 124,891 | 23,029 | 18,979 |
| Map resolution (Å) | 3.6 | 4.3 | 4.9 |
| FSC threshold | 0.143 | 0.143 | 0.143 |
| Map sharpening B factor (Å$^2$) | -80 | -40 | -40 |

*Table S2* ***Interface area between adjacent protomers of the Nipah virus (NiV), Parainfluenza virus 5 (PIV5) and Measles virus (MeV).*** *The buried surface area was calculated for the helical assembly of each virus.*

| | NiV (This study) | PIV5 (PDB:4xjn) | MeV (PDB:6h5q) |
|---|---|---|---|
| Interface area (A$^2$) | 2998 (100%) | 2918 (100%) | 2931 (100%) |
| Polar interface area (Å$^2$) | 2011 (67%) | 1931 (66%) | 1812 (62%) |
| Nonpolar interface area (Å$^2$) | 986 (33%) | 987 (34%) | 1119 (38%) |

*Table S3* ***Domain movements in the NiV N protein around the hinge area associated with RNA binding.*** *Rotational and translational values were derived from comparison of the RNA-free (pdb:4co6) and RNA-bound states. All values were estimated as described in Methods.*

| | |
|---|---|
| Rotation Angle (°) | 27.9 |
| Translation (Å) | 1.8 |
| Closure (%) | 46.8 |
| Hinge region residues | 263-265<br>303-304<br>317-321 |

*Table S4 RMSD between the RNA-bound (this study) and RNA-free (pdb:4co6) NiV N protein calculated for Cα atoms.*

| Region | RMSD (Å) |
|---|---|
| Whole protein (residue 32-369) | 3.3 |
| N-terminal Ncore (residue 32-258) | 1.7 |
| C-terminal Ncore (residue 286-369) | 1.7 |
